# ModularImageAnalysis (MIA): Assembly of modularised image and object analysis workflows in ImageJ

**DOI:** 10.1101/2023.06.12.544614

**Authors:** Stephen J. Cross, Jordan D. J. R. Fisher, Mark A. Jepson

## Abstract

ModularImageAnalysis (MIA) is an ImageJ plugin providing a code-free graphical environment in which complex automated analysis workflows can be constructed and distributed. The near 200 included modules cover all stages of a typical analysis workflow, from image loading through image processing, object detection, extraction of measurements, measurement-based filtering, visualisation, and data exporting. MIA provides out-of-the-box compatibility with many advanced image processing plugins for ImageJ including Bio-Formats, DeepImageJ, MorphoLibJ and TrackMate, allowing these tools and their outputs to be directly incorporated into analysis workflows. By default, modules support spatially calibrated 5D images, meaning measurements can be acquired in both pixel and calibrated units. A hierarchical object relationship model allows for both parent-child (one-to-many) and partner (many-to-many) relationships to be established. These relationships underpin MIA’s ability to track objects through time, represent complex spatial relationships (e.g. skeletons) and measure object distributions (e.g. count puncta per cell). MIA features dual graphical interfaces: the “editing view” offers access to the full list of modules and parameters in the workflow, while the simplified “processing view” can be configured to display only a focused subset of controls. All workflows are batch-enabled by default, with image files within a specified folder being processed automatically and exported to a single spreadsheet. Beyond the included modules, functionality can be extended both internally, through integration with the ImageJ scripting interface, and externally, by developing third-party Java modules that extend the core MIA framework. Here we describe the design and functionality of MIA in the context of a series of real-world example analyses.

## Introduction

In the field of bioimage analysis, few tools have gained the widespread popularity of ImageJ and commonly used variants such as Fiji ^1–3^. This prolific uptake can in a large part be attributed to ImageJ’s open-source nature and easy extensibility via macros and plugins. Over the past 25 years, the catalogue of available plugins has endowed the ImageJ ecosystem with a vast and comprehensive functionality, covering features as diverse as object detection and tracking ^4,5^, tiled image stitching and drift correction ^6–8^, morphometric analysis ^9^, 3D rendering ^10,11^, and recently, integration with deep-learning frameworks ^12,13^. By providing a singular environment in which these tools can work together, ImageJ very much becomes more than a sum of its parts.

Despite ImageJ’s versatility, construction and distribution of reproducible and automated workflows remains challenging, as users generally interact with plugins via either graphical user interfaces or programmatically with macros and scripts. While graphical user interfaces may offer a user-friendly manner of operation, manually working through a series of steps can be prohibitive to analysis of large datasets. Workflow reproducibility under this environment is also of concern, as it relies solely on accurate documentation of every step down to the tiniest detail ^14^. Conversely, the need for, at minimum, a rudimentary understanding of programming in order to assemble macros and scripts can be off-putting to newcomers. ImageJ’s integrated macro recorder certainly lowers the barrier to assembly of such workflows; however, macros created in this manner often require refinement and the code needed to run some plugins cannot be fully captured via the recorder.

To make bioimage analysis with ImageJ more accessible to a wider community, we present ModularImageAnalysis (MIA), an ImageJ plugin for code-free assembly of complex image and object analysis workflows. With MIA, images and objects are passed between self-contained modules, each of which performs a distinct operation in an analysis workflow, such as image filtering, object detection, measurement of object properties and visualisation of results. By utilising a single object format, objects detected in one module can be immediately passed to downstream modules without the need for conversion or translation. At present, MIA offers approximately 200 modules, many of which integrate advanced image processing plugins for ImageJ such as Bio-Formats ^15^, TrackMate ^4^, Trainable Weka Segmentation ^16^, MorphoLibJ ^9^ and DeepImageJ ^12^. Workflows are assembled in a modularised manner which will be familiar to users of similar tools such as CellProfiler ^17^, Icy ^18^ and KNIME ^19^, whilst adding several novel features not found elsewhere. MIA also addresses the issue of workflow reproducibility and documentation, by storing workflows in an easy to distribute text-based format, which has been designed with the FAIR Data Principles in mind ^20^.

Here we will describe the core concepts behind MIA, such as its dual user interfaces, one for workflow creators and the other for end users, its compatibility with a wide range of multidimensional image formats and the strategies employed to ensure optimal computational performance. This will be presented in the context of a range of real-world examples spanning multiple imaging modalities.

### MIA workflow structures

Workflows in MIA are comprised of a sequence of modules, with each module handling a specific task, such as image loading, object detection or calculation of measurements. Each module can output items, including images, objects and measurements, to a common workspace. The items in this workspace are assigned user-defined names, allowing subsequent modules in the workflow to access them. The workspace is unique to a single analysis run; as such, when processing multiple images (batch mode), multiple non-interacting workspaces are created.

A simple example workflow is shown in Figure 1 and depicts the segmentation of cell nuclei from a fluorescence microscopy image (workflow file adapted from example at ^21^). In this example, the first module (“Load image”) reads an image from file and stores it in the workspace with the name “Raw”. This image is then accessed by the next module (“Apply threshold”), which applies an automatically calculated intensity threshold and stores the resulting binarised image in the workspace as a new image called “Binary”. In this instance, the original “Raw” image was unaltered by execution of the “Apply threshold” module; however, there may be instances where the input image is no longer required in its original form, as is the case with the “Fill holes” module in Figure 1. Here, the input image can simply be updated within the workspace, eliminating unnecessary memory usage. This is especially useful when dealing with large, multidimensional image stacks.

**Figure 1.**
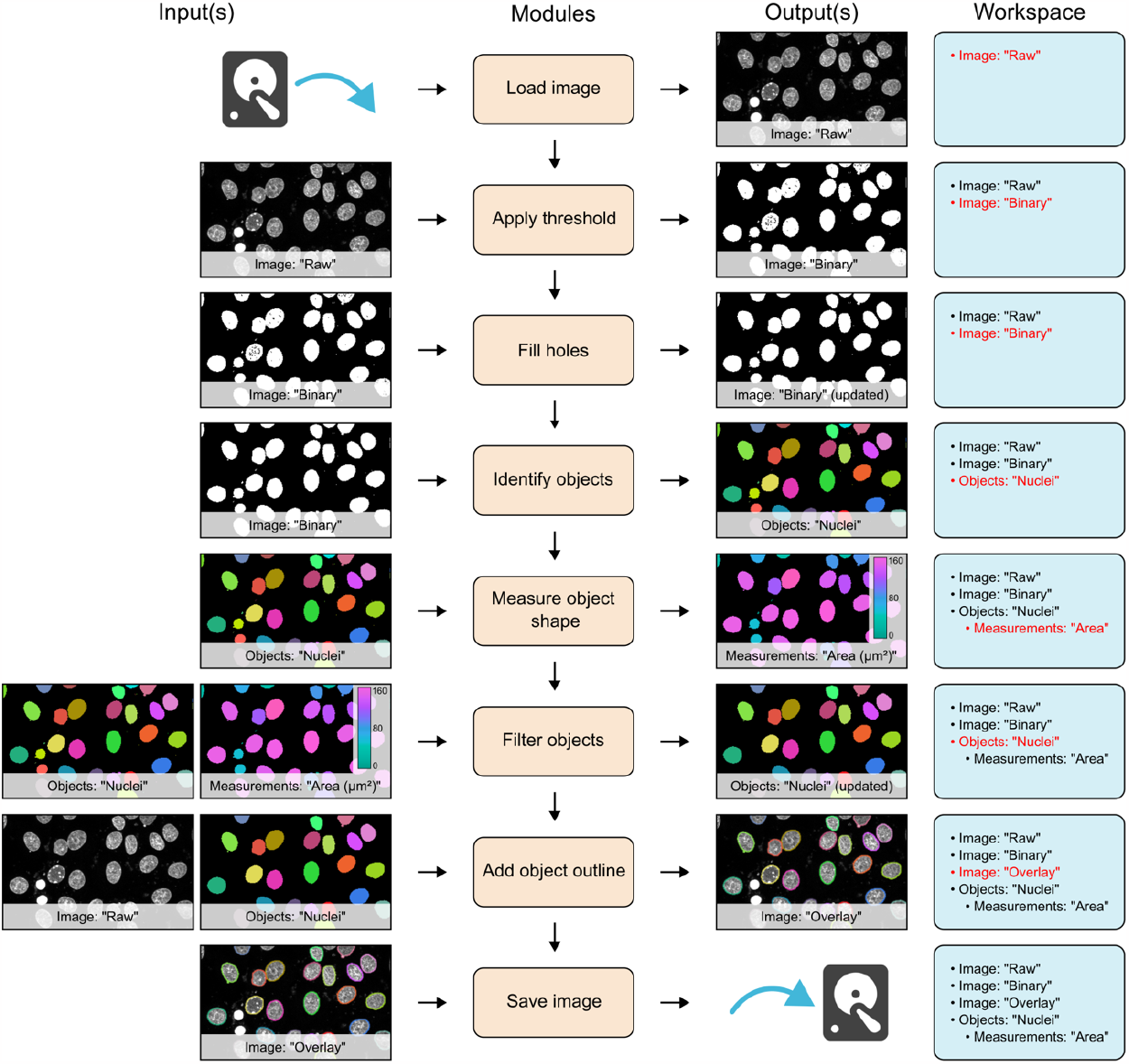
Schematic of a simple workflow for segmentation of cell nuclei. Workflow is depicted with each module on a separate row. To the left of the module list are the input images, objects and measurements for that module. Likewise, outputs are shown to the right of the module list. On the far right is a list of all images, objects and modules available in the workspace at each stage. New or updated items in the workspace are highlighted in red.

In addition to images, modules can also read and write objects to the workspace. In the workflow shown in Figure 1, the “Identify objects” module applies connected components labelling to identify individual objects in the “Binary” image, then stores these as the object set, “Nuclei”. Both images and objects can have measurements associated with them. Here, the “Measure object shape” module calculates the spatial area of the objects stored as “Nuclei” and the subsequent “Filter objects” module uses these values, along with a user-defined threshold, to exclude any small objects likely arising from noise in the original image. A suite of visualisation modules can be used to, amongst other operations, add ImageJ-compatible overlays to images. In Figure 1, random colours are assigned to the identified objects in order to distinguish them and the outlines of these objects are added to the “Raw” image, which is then written to file using the “Save image” module.

In MIA, workflows are batch processing-enabled by default. By setting the input file path to a folder, rather than a single file, MIA will detect all valid files within that folder (and sub-folders) and process them as independent “jobs”. An arbitrary number of filters based on file, folder and series names can be applied to provide selective processing of files during batch operation. Furthermore, through its fundamental integration of the Bio-Formats library ^15^, MIA is capable of reading a wide array of open and proprietary imaging formats. This includes multi-series files (for example, Leica LIF and Olympus VSI), from which all, or a subset of, series can be automatically processed as a batch. MIA gives the flexibility to store results according to preferred experimental design; chiefly, all output measurements are optional, simple measurements statistics (mean, minimum, maximum, etc.) can be calculated on a per-image basis and the results from a single batch run can be combined into a single Excel XLSX file or stored individually.

To date, MIA has nearly 200 modules; these can be broadly placed into the nine categories shown in Figure 2. Many of these modules interact with popular ImageJ plugins, allowing these tools to be seamlessly integrated into automated workflows without the need for additional scripts or data manipulation. This interoperability is facilitated by MIA’s use of a single object format, which stores pixel coordinates for each object along with measurements and object-object relationships.

**Figure 2.**
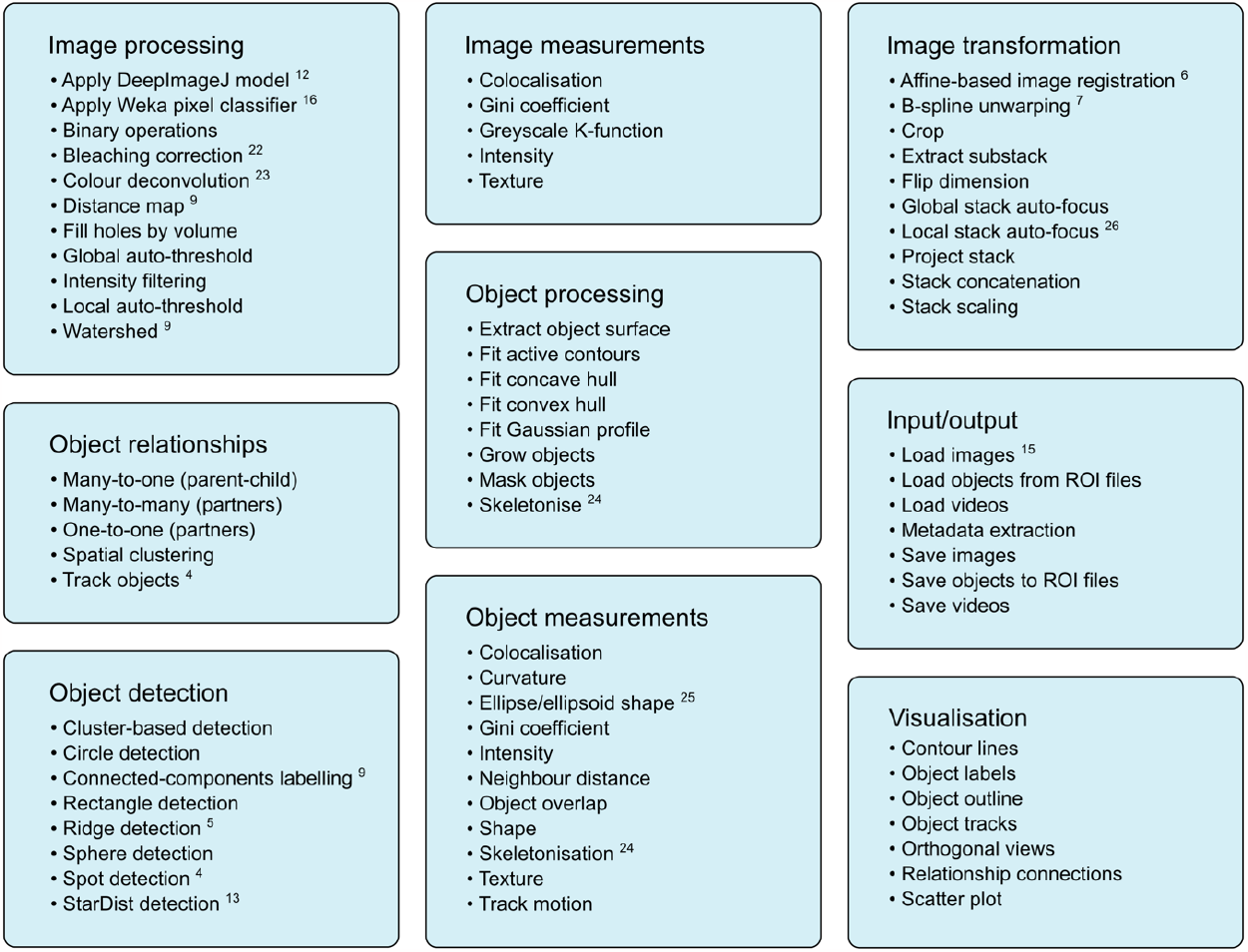
Example modules showcasing the range of functionality within MIA. The nine main module categories with examples for each. Many modules interact and utilise existing ImageJ plugins such as DeepImageJ ^12^, Trainable Weka Segmentation ^16^, Bleach Correction ^22^, Colour Deconvolution ^23^, MorphoLibJ ^9^, TrackMate ^4^, Ridge Detection ^5^, StarDist ^13^, Analyze Skeletons ^24^, BoneJ ^25^, TrakEM2 ^6^, bUnwarpJ ^7^, Stack Focuser ^26^ and Bio-Formats ^15^.

Workflows are stored as text files (with the .mia extension), thus allowing for easy sharing and reuse as well as addressing the issue of workflow reproducibility. Moreover, the full workflow configuration is also stored in each exported XLSX file, meaning workflows can be recalled in cases where the original workflow .mia file has been lost or altered.

### User interface

MIA offers two user interfaces, a fully featured “Editing view” and a simplified “Processing view” (Figure 3). For workflow creators, “Editing view” provides an environment in which to assemble modules into workflows. Modules are arranged into an execution-ordered list, with parameters for each module displayed upon clicking a module’s name.

**Figure 3.**
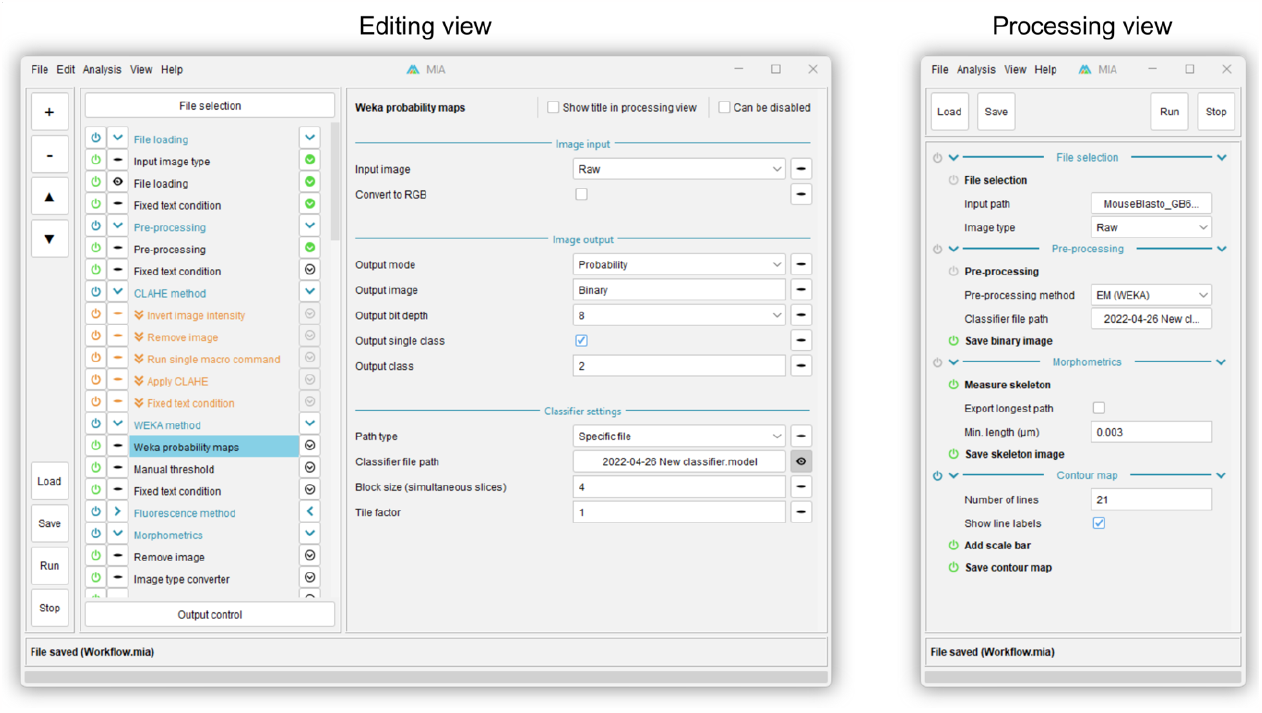
Screenshots of the MIA user interface. Editing view (left) is designed for creating and editing workflows and offers access to all modules and parameters in a workflow. Processing view (right) is intended for running pre-assembled workflows and is typically configured (via Editing view) to display a small sub-set of parameters required to run an analysis on new images.

To facilitate easy workflow design and troubleshooting, modules can be executed one-by-one using arrow buttons to the right of each module name. The output from a module (for example, an image showing detected objects) can be visualised by enabling the eye button to the left of that module’s name. Modules can also be enabled and disabled using power icons to the left of each module name. Since workflows are reactive, disabling a module will automatically disable any downstream modules which relied on the outputs of the disabled module. Likewise, a collection of “workflow handling” modules can be used to skip modules or terminate workflow execution entirely based on user-defined conditions (skipped modules shown in orange in Figure 3).

The “Processing view” is designed for end users and day-to-day running of workflows; this view displays the subset of parameters required to run a workflow on new images, such as input file/folder locations. It may also be configured to show controls for fine-tuning workflows, for example object size filters or data exporting options. By making these controls accessible in a simplified view, non-expert users can configure key parts of the workflow without needing to delve into the potentially complex workings of the full workflow otherwise available via “Editing view”. The controls visible in “Processing view” can be selected in “Editing view” by enabling the eye button to the right of each parameter (for example, the “Classifier file path” parameter shown in Figure 3).

### Object relationships

Individual objects in MIA are stored in the three spatial dimensions (XYZ); however, there are instances where it’s necessary to consider their existence along additional axes. Examples include tracking objects across multiple timeframes or cases where the same object is identified in different channels of an image stack. To capture these relationships, MIA supports both parent-child (one-to-many) and partner (many-to-many) relationships. In the parent-child relationship example shown in Figure 4, nuclei are first detected in individual time frames, then tracked between these frames using the “Track objects” module, which utilises TrackMate’s Jaqaman linker algorithm ^4,27^. Each track is stored as a new “Track” object, which itself contains no spatial (coordinate) information; instead, each “Track” object acts as a parent to the child “Nuclei” objects in that track. With objects tracked between frames it becomes possible to measure temporal characteristics, such as velocity, directionality and total path length using the “Measure track motion” module.

**Figure 4.**
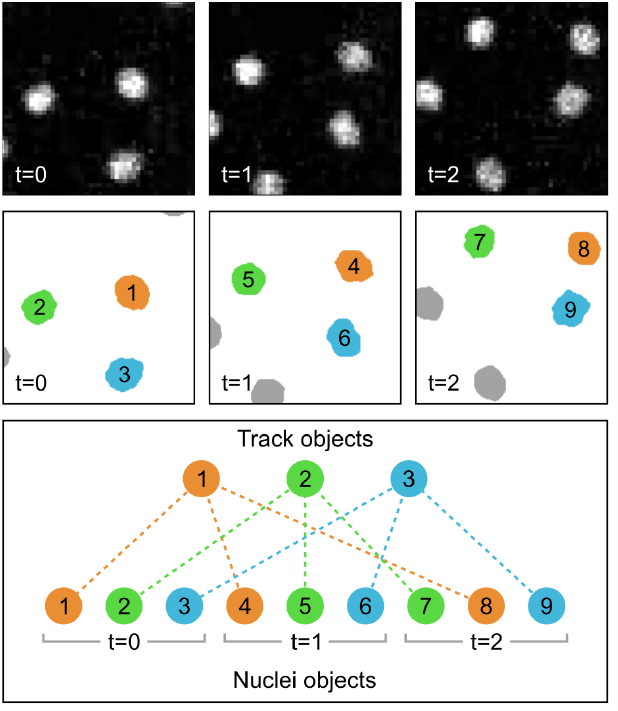
Schematic diagram showing parent-child object relationships in the context of object tracking. In this example, nuclei from three frames of a fluorescent image timeseries (top row) are detected and stored as “Nuclei” objects, each of which is given a unique ID number (middle row). These individual “Nuclei” objects are tracked across each frame and assigned as children to newly created “Track” objects. These “Track” objects simply act as linking objects and themselves contain no spatial (coordinate) information. The relationships are depicted by both colour-coding and as dashed lines between the track and nuclei objects (bottom row). Fluorescent images are of yolk syncytial layer (YSL) nuclear movements cropped from ^28^.

Importantly, the coordinate-less “linking” behaviour of tracks isn’t the sole purpose of parent-child relationships. In an alternative example, a previously detected parent “Cell” object could be subdivided into “Interior” and “Exterior” child objects based on the distance of coordinates from the surface of that cell. Here, the pixel intensity of a separate fluorescence channel could be measured for the “Interior” and “Exterior” objects, then used to calculate an interior:exterior signal ratio for each parent “Cell”.

Objects can also be engaged in multiple parent-child relationships. The implication of this being that while child objects can only have one parent of a given class, they may have multiple parents across many different classes. MIA also supports extended relationships, effectively enabling grandparent-grandchild relationships and beyond. By combining the two aforementioned examples of object tracking and object subdivision we could have a scenario where the tracked objects (for example, cells) are subdivided at each timepoint into interior and exterior objects. These interiors and exteriors would then be grandchildren of the track objects, giving the opportunity to measure the average interior:exterior intensity ratio across all timepoints.

Partner relationships are used in cases where objects in both associated classes may be linked to multiple other objects. Figure 5 shows one such example, with grain boundaries in SEM images being represented as “Edge” and “Junction” objects. Here, edges will be related to up to two junctions and each junction will have multiple associated edges. This allows complex relationships to be captured. In the given example, it would be possible to remove edges connected to only one junction (i.e. branches of the skeleton). As with parent-child relationships, objects can be engaged in partner relationships with multiple object classes.

**Figure 5.**
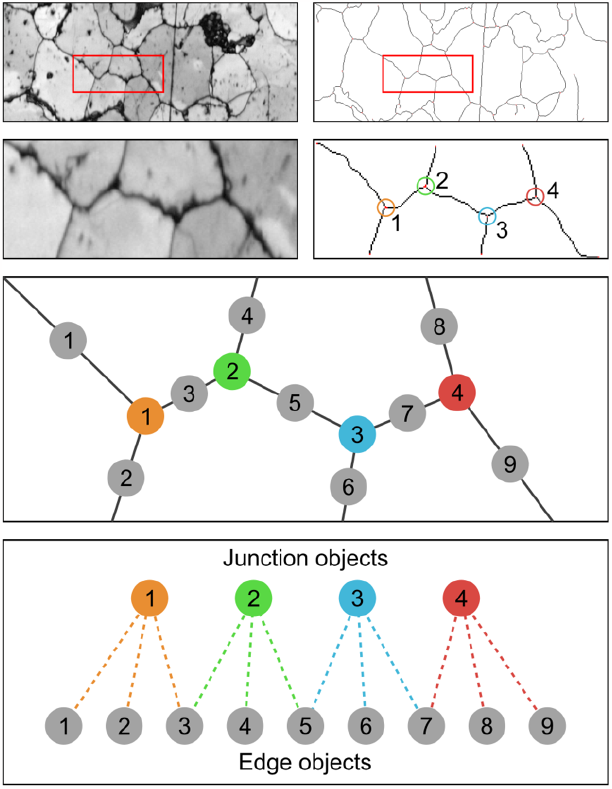
Schematic diagram showing partner object relationships applied to grain boundary analysis of a deformed quartzite. Binarised grain boundaries are skeletonised, with the skeleton fragments stored as either “Junction” (shown with assorted colours) or “Edge” (shown in grey) objects. These objects are assigned partner relationships, where each “Junction” or “Edge” can be linked to multiple instances of the other object. Raw images are band contrast maps from electron backscatter diffraction (EBSD) analysis ^29^. Image courtesy of A. Cross (Woods Hole Oceanographic Institution, Massachusetts, USA).

### Memory efficient coordinate storage

Object handling necessitates the storage of pixel coordinates in a form that can be easily accessed by downstream modules. In the case of small objects (for example, foci detected by spot detecting modules), the most practical solution is simply to record these as lists of XYZ pixel locations (henceforth referred to as “pointlists”); however, for storage of large regions, this approach can be inefficient and memory-limiting. In commonly occurring instances where large regions are comprised of significant contiguous and hole-free areas, efficient coordinate storage can be achieved using quadtrees ^30^. Quadtrees recursively subdivide an area into four nodes (quadrants), with subdivision ending once a node entirely contains either foreground (object) or background coordinates. As such, large contiguous regions can be summarised by relatively few nodes and in extreme cases reduce memory requirements by orders of magnitude compared to the equivalent pointlist structure (Figure 6).

**Figure 6.**
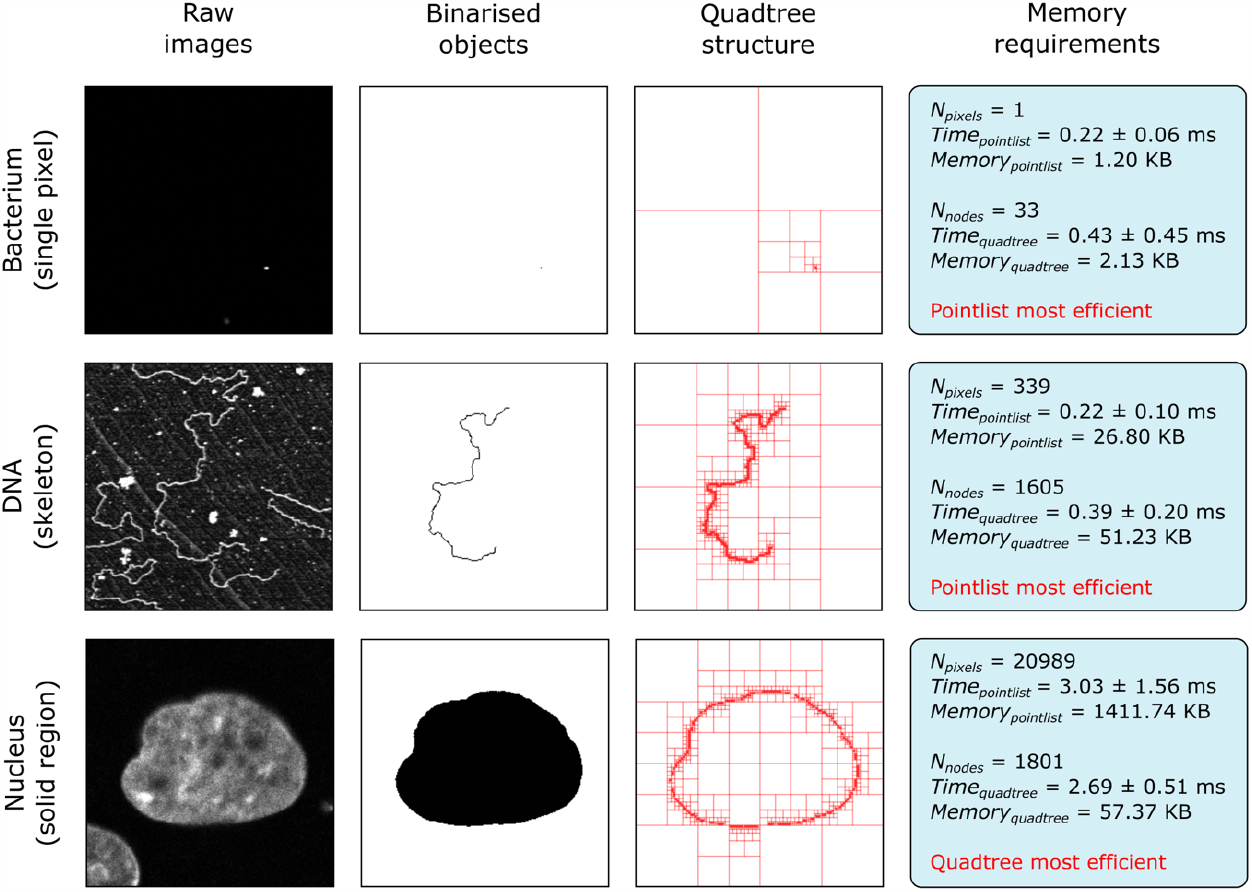
Comparison of coordinate storage methods. Performance of storing object coordinates as pointlists and quadtrees for three different object morphologies. Shown for each example are the raw image, binarised form of the object and quadtree representation. The two storage methods are evaluated on the time taken to create the coordinate store and the memory it occupies. Time taken is reported as the average of 30 measurements with standard deviation. For each sample, the most efficient memory storage method is highlighted in red. All three raw images have a resolution of 256 × 256 px^2^.

Figure 6 shows a performance comparison between pointlists and quadtrees when storing three example objects with differing morphologies. Here, the “Memory” metric relates to the memory required to store the object and the “Time” metric is the time taken to convert the binarised image to the final data structure (reported as the average of 30 measurements with standard deviation). For storage of a single pixel (“Bacterium” sample in Figure 6), pointlists offer both a memory and speed advantage, due to the need of quadtrees to subdivide the image region down to the single pixel level. When storing single pixel objects, this performance difference will increase with image size, as more subdivisions are required. Likewise, for skeletonised objects (“DNA” sample in Figure 6), the necessary subdivision to the pixel level yields similar performance. Quadtrees begin to outperform pointlists for solid regions (“Nucleus” sample in Figure 6), where the large number of coordinates in the centre of the object can be simplified by relatively few nodes. In such samples, subdivision to the pixel level is only required on the edge of the object; therefore, the number of nodes used can be a fraction of the total pixel count.

MIA also includes an octree storage option, with octrees behaving as the 3D counterpart to quadtrees, albeit with image stack subdivision into octants. Octrees perform best with image stacks acquired using isotropic spatial resolution, as is typically the case with micro-CT. For modules outputting objects where a specific coordinate storage method will always be most efficient (for example, pointlists for spot detection and skeletonisation), that method is used exclusively. Conversely, general purpose object detection modules offer the user the choice of object storage method. All coordinate storage approaches provide the same core functionality (by extending the same Java class), so their use is interchangeable with no difference to the end-user beyond memory performance.

### Extensibility

Beyond the approximately 200 included modules, MIA’s functionality can be extended both internally, through integration with Fiji’s scripting interface, and externally, by developing new modules that extend the core MIA framework. At the simplest level, the “Run single command” module, allows individual ImageJ macro commands of the form run(command, parameters) to be applied to images in the workspace. For more complex applications, the “Run script” module can be used to run scripts with full access to the current analysis workspace. This module supports any language compatible with ImageJ’s scripting interface, including Beanshell, Groovy, Javascript and Jython. Scripts have the ability to create new images and object collections, add image and object-associated measurements and assign new object relationships. Using MIA’s application programming interface (API), these actions can each be achieved from a script with a couple of commands. An example where such functionality may be useful is in calculating custom measurements for objects detected earlier in a workflow. Finally, custom code that may be routinely used across multiple workflows can be packaged as entirely new modules, which are distributed as MIA plugins. Since MIA uses SciJava’s “Plugin” interface ^31^, any Java classes extending MIA’s “Module” class will be automatically available for inclusion in workflows in the same manner as standard integrated modules. Indeed, MIA plugins already exist for operations that require large additional libraries that are impractical to bundle with the main MIA distribution. Examples include the “Load videos” module, which requires platform-specific binaries, and “Fit alpha surface”, which requires the MATLAB Compiler Runtime to be installed.

In addition to incorporating new functionality into MIA workflows via scripting, MIA offers a suite of macro commands allowing the functionality of MIA to be incorporated into macro-based workflows. These commands provide a route to run workflows directly from standard ImageJ macros, that is, outside of the MIA graphical user interface. Furthermore, and irrespective of whether the workflow was initially run via the MIA interface or from a macro, these commands give macros access to the individual images and objects within each analysis workspace, as well as to all their associated measurements and relationships.

## Discussion

To date, MIA has been used in a wide variety of bioimage analyses, including: 3D detection of fluorescent puncta ^32^, quantification of immune cell behaviour in response to cancer ^33^, morphometric analysis of extracellular DNA ^34^, analysis of cell morphology in epithelial tissue ^35^, detection of cellular membrane tubules ^36^ and 3D assessment of zebrafish vertebrae from micro-CT image stacks ^37^.

Through these examples, we can observe the flexibility and versatility of MIA; by providing an environment in which the strengths present in the extensive catalogue of ImageJ plugins can be combined, it’s possible to create workflows with comprehensive functionality. Furthermore, MIA also addresses the important ongoing challenge of image analysis workflow reproducibility as well as their distribution.

As with the ImageJ ecosystem itself, MIA is under constant development in response to advancements in the field of image processing and analysis. Modules supporting both new ImageJ plugins and custom functionality are added in response to the needs of users and ongoing analyses. In particular, recent developments have seen the inclusion of modules implementing popular deep learning plugins such as DeepImageJ ^12^ and StarDist ^13^.

More fundamentally, the increasing prevalence of imaging modalities generating very large image volumes (taken here to be 10s of GB and greater), such as lightsheet and serial block face SEM, are being addressed by changes to MIA’s image handling and storage framework. Currently, MIA stores images using ImageJ’s ImagePlus format ^1^, which relies on first loading images into RAM. This provides fast response times when accessing pixel information, but places limits on the size of image volume that can be processed at one time. Ongoing development sees a transition from ImagePlus to the newer ImgLib2 format which also powers ImageJ2 ^2,38^. With ImgLib2’s disk-cached image formats, when an image would exceed the available memory, pixel data can be read directly from a computer’s filesystem without the need to preload it into RAM. Similarly, new images with sizes exceeding the available memory can be written to a new disk cache, effectively removing the upper limit on the size of image volumes, albeit with a performance penalty based on the speed of the available storage. For images capable of fitting in the available RAM, performance should be equivalent to the original ImagePlus-based system.

Through its ongoing development and with the built-in capacity for extension via user-driven scripts, MIA has the potential to be employed for an increasingly diverse range of bioimage analyses. Furthermore, by offering a graphical environment in which to assemble workflows, MIA democratises the use of ImageJ for state of the art analysis in an era where the acquisition and handling of images is becoming a fundamental part of many experimental designs.

## Data availability

Source code is available at https://github.com/mianalysis/mia. Full documentation, including tutorial videos and example workflows, can be accessed from https://mianalysis.github.io.

## Acknowledgements

The authors would like to thank all users of the Wolfson Bioimaging Facility (Bristol) who have worked with and provided invaluable feedback on MIA. Special thanks to Dominic Alibhai for many years of testing and feature suggestions and to Gemma Fisher for preparing automated tests of many modules. Finally, thanks to Ana Stojiljković, Alin Achim and Andrew Cross for manuscript proofreading and feedback. Stephen Cross was part funded by the Elizabeth Blackwell Institute, through its Wellcome Trust ISSF Award.

## References

1. Schneider, C.A., Rasband, W.S., and Eliceiri, K.W. (2012) NIH Image to ImageJ: 25 years of image analysis. Nat. Methods, 9 (7), 671–675.

2. Rueden, C.T., Schindelin, J., Hiner, M.C., DeZonia, B.E., Walter, A.E., Arena, E.T., and Eliceiri, K.W. (2017) ImageJ2: ImageJ for the next generation of scientific image data. BMC Bioinformatics, 18 (1), 529.

3. Schindelin, J., Arganda-Carreras, I., Frise, E., Kaynig, V., Longair, M., Pietzsch, T., Preibisch, S., Rueden, C., Saalfeld, S., Schmid, B., Tinevez, J.-Y., White, D.J., Hartenstein, V., Eliceiri, K., Tomancak, P., and Cardona, A. (2012) Fiji: an open-source platform for biological-image analysis. Nat. Methods, 9 (7), 676– 682.

4. Tinevez, J.-Y., Perry, N., Schindelin, J., Hoopes, G.M., Reynolds, G.D., Laplantine, E., Bednarek, S.Y., Shorte, S.L., and Eliceiri, K.W. (2017) TrackMate: An open and extensible platform for single-particle tracking. Methods, 115, 80– 90.

5. Wagner, T., Hiner, M., and Xraynaud (2017) thorstenwagner/ij-ridgedetection: Ridge Detection 1.4.0.

6. Cardona, A., Saalfeld, S., Schindelin, J., Arganda-Carreras, I., Preibisch, S., Longair, M., Tomancak, P., Hartenstein, V., and Douglas, R.J. (2012) TrakEM2 Software for Neural Circuit Reconstruction. PLoS ONE, 7 (6), e38011.

7. Arganda-Carreras, I., Sorzano, C.O.S., Marabini, R., Carazo, J.M., Ortiz-de-Solorzano, C., and Kybic, J. (2006) Consistent and Elastic Registration of Histological Sections Using Vector-Spline Regularization, in Computer Vision Approaches to Medical Image Analysis, vol. 4241, Springer Berlin Heidelberg, Berlin, Heidelberg, pp.85–95.

8. Preibisch, S., Saalfeld, S., and Tomancak, P. (2009) Globally optimal stitching of tiled 3D microscopic image acquisitions. Bioinformatics, 25 (11), 1463–1465.

9. Legland, D., Arganda-Carreras, I., and Andrey, P. (2016) MorphoLibJ: integrated library and plugins for mathematical morphology with ImageJ. Bioinformatics, 32 (22), 3532–3534.

10. Schmid, B., Tripal, P., Fraaß, T., Kersten, C., Ruder, B., Grüneboom, A., Huisken, J., and Palmisano, R. (2019) 3Dscript: animating 3D/4D microscopy data using a natural-language-based syntax. Nat. Methods, 16 (4), 278–280.

11. Schmid, B., Schindelin, J., Cardona, A., Longair, M., and Heisenberg, M. (2010) A high-level 3D visualization API for Java and ImageJ. BMC Bioinformatics, 11 (1), 274.

12. Gómez-de-Mariscal, E., García-López-de-Haro, C., Ouyang, W., Donati, L., Lundberg, E., Unser, M., Muñoz-Barrutia, A., and Sage, D. (2021) DeepImageJ: A user-friendly environment to run deep learning models in ImageJ. Nat. Methods, 18 (10), 1192–1195.

13. Schmidt, U., Weigert, M., Broaddus, C., and Myers, G. (2018) Cell Detection with Star-Convex Polygons, in Medical Image Computing and Computer Assisted Intervention – MICCAI 2018, vol. 11071, Springer International Publishing, Cham, pp.265–273.

14. Miura, K., and Nørrelykke, S.F. (2021) Reproducible image handling and analysis. EMBO J., 40 (3), e105889.

15. Linkert, M., Rueden, C.T., Allan, C., Burel, J.-M., Moore, W., Patterson, A., Loranger, B., Moore, J., Neves, C., Macdonald, D., Tarkowska, A., Sticco, C., Hill, E., Rossner, M., Eliceiri, K.W., and Swedlow, J.R. (2010) Metadata matters: access to image data in the real world. J. Cell Biol., 189 (5), 777–782.

16. Arganda-Carreras, I., Kaynig, V., Rueden, C., Eliceiri, K.W., Schindelin, J., Cardona, A., and Sebastian Seung, H. (2017) Trainable Weka Segmentation: a machine learning tool for microscopy pixel classification. Bioinformatics, 33 (15), 2424–2426.

17. Stirling, D.R., Swain-Bowden, M.J., Lucas, A.M., Carpenter, A.E., Cimini, B.A., and Goodman, A. (2021) CellProfiler 4: improvements in speed, utility and usability. BMC Bioinformatics, 22 (1), 433.

18. De Chaumont, F., Dallongeville, S., Chenouard, N., Hervé, N., Pop, S., Provoost, T., Meas-Yedid, V., Pankajakshan, P., Lecomte, T., Le Montagner, Y., Lagache, T., Dufour, A., and Olivo-Marin, J.-C. (2012) Icy: an open bioimage informatics platform for extended reproducible research. Nat. Methods, 9 (7), 690–696.

19. Berthold, M.R., Cebron, N., Dill, F., Gabriel, T.R., Kötter, T., Meinl, T., Ohl, P., Sieb, C., Thiel, K., and Wiswedel, B. (2008) KNIME: The Konstanz Information Miner, in Data Analysis, Machine Learning and Applications (eds.Preisach, C., Burkhardt, H., Schmidt-Thieme, L., and Decker, R.), Springer Berlin Heidelberg, Berlin, Heidelberg, pp.319–326.

20. Wilkinson, M.D., Dumontier, M., Aalbersberg, Ij.J., Appleton, G., Axton, M., Baak, A., Blomberg, N., Boiten, J.-W., Da Silva Santos, L.B., Bourne, P.E., Bouwman, J., Brookes, A.J., Clark, T., Crosas, M., Dillo, I., Dumon, O., Edmunds, S., Evelo, C.T., Finkers, R., Gonzalez-Beltran, A., Gray, A.J.G., Groth, P., Goble, C., Grethe, J.S., Heringa, J., ‘T Hoen, P.A.C., Hooft, R., Kuhn, T., Kok, R., Kok, J., Lusher, S.J., Martone, M.E., Mons, A., Packer, A.L., Persson, B., Rocca-Serra, P., Roos, M., Van Schaik, R., Sansone, S.-A., Schultes, E., Sengstag, T., Slater, T., Strawn, G., Swertz, M.A., Thompson, M., Van Der Lei, J., Van Mulligen, E., Velterop, J., Waagmeester, A., Wittenburg, P., Wolstencroft, K., Zhao, J., and Mons, B. (2016) The FAIR Guiding Principles for scientific data management and stewardship. Sci. Data, 3 (1), 160018.

21. Cross, Stephen John (2023) MIA Examples (version 1.0.1).

22. Miura, K. (2020) Bleach correction ImageJ plugin for compensating the photobleaching of time-lapse sequences. F1000Research, 9, 1494.

23. Landini, G., Martinelli, G., and Piccinini, F. (2021) Colour deconvolution: stain unmixing in histological imaging. Bioinformatics, 37 (10), 1485–1487.

24. Arganda-Carreras, I., Fernández-González, R., Muñoz-Barrutia, A., and Ortiz-De-Solorzano, C. (2010) 3D reconstruction of histological sections: Application to mammary gland tissue. Microsc. Res. Tech., 73 (11), 1019–1029.

25. Domander, R., Felder, A.A., and Doube, M. (2021) BoneJ2 – refactoring established research software. Wellcome Open Res., 6, 37.

26. Umorin, M. (2006) Stack Focuser.

27. Jaqaman, K., Loerke, D., Mettlen, M., Kuwata, H., Grinstein, S., Schmid, S.L., and Danuser, G. (2008) Robust single-particle tracking in live-cell time-lapse sequences. Nat. Methods, 5 (8), 695–702.

28. D’Amico, L. (2011) IL_11813, Danio rerio, yolk syncytial layer cell.

29. Cross, A.J., Prior, D.J., Stipp, M., and Kidder, S. (2017) The recrystallized grain size piezometer for quartz: An EBSD-based calibration: EBSD-Based Quartz Grain Size Piezometer. Geophys. Res. Lett., 44 (13), 6667–6674.

30. Finkel, R.A., and Bentley, J.L. (1974) Quad trees a data structure for retrieval on composite keys. Acta Inform., 4 (1), 1–9.

31. Rueden, C., Schindelin, J., Hiner, M., and Eliceiri, K. (2021) SciJava Common.

32. McCaughey, J., Stevenson, N.L., Cross, S., and Stephens, D.J. (2019) ER-to-Golgi trafficking of procollagen in the absence of large carriers. J. Cell Biol., 218 (3), 929–948.

33. López-Cuevas, P., Cross, S.J., and Martin, P. (2021) Modulating the Inflammatory Response to Wounds and Cancer Through Infection. Front. Cell Dev. Biol., 9, 676193.

34. Serrage, H.J., FitzGibbon, L., Alibhai, D., Cross, S., Rostami, N., Jack, A.A., Lawler, C.R.E., Jakubovics, N.S., Jepson, M.A., and Nobbs, A.H. (2022) Quantification of Extracellular DNA Network Abundance and Architecture within Streptococcus gordonii Biofilms Reveals Modulatory Factors. Appl. Environ. Microbiol., 88 (13), e00698–22.

35. Olenik, M., Turley, J., Cross, S., Weavers, H., Martin, P., Chenchiah, I.V., and Liverpool, T.B. (2023) Fluctuations of cell geometry and their nonequilibrium thermodynamics in living epithelial tissue. Phys. Rev. E, 107 (1), 014403.

36. Stan, G.F., Shoemark, D.K., Alibhai, D., and Hanley, J.G. (2022) Ca2+ Regulates Dimerization of the BAR Domain Protein PICK1 and Consequent Membrane Curvature. Front. Mol. Neurosci., 15, 893739.

37. Kague, E., Turci, F., Newman, E., Yang, Y., Brown, K.R., Aglan, M.S., Otaify, G.A., Temtamy, S.A., Ruiz-Perez, V.L., Cross, S., Royall, C.P., Witten, P.E., and Hammond, C.L. (2021) 3D assessment of intervertebral disc degeneration in zebrafish identifies changes in bone density that prime disc disease. Bone Res., 9 (1), 39.

38. Pietzsch, T., Preibisch, S., Tomančák, P., and Saalfeld, S. (2012) ImgLib2— generic image processing in Java. Bioinformatics, 28 (22), 3009–3011.

